# Proximity of pre-mRNA 3′ end processing and transcription termination predicts enhanced gene expression

**DOI:** 10.64898/2025.12.02.691783

**Authors:** Agata Stępień, Deepshika Pulimamidi, Martyna Plens-Gałąska, Maxime Lalonde, Magda Kopczyńska, Hiroshi Kimura, Shazia Irshad, Michał R. Gdula, Stephan Hamperl, Kinga Kamieniarz-Gdula

**Affiliations:** Center for Advanced Technologies, Adam Mickiewicz University, 61-614, Poznań, Poland; Department of Molecular and Cellular Biology, Institute of Molecular Biology and Biotechnology, Faculty of Biology, Adam Mickiewicz University, 61-614, Poznań, Poland; Chromosome Dynamics and Genome Stability, Institute of Epigenetics and Stem Cells, Helmholtz Zentrum München, 81377, Munich, Germany; Cell Biology Centre, Institute of Innovative Research, Tokyo Institute of Technology, 226-8503, Yokohama, Japan; Nuffield Department of Clinical Laboratory Sciences, Radcliffe Department of Medicine, University of Oxford, OX3 9DU, Oxford, UK; Department of Gene Expression, Institute of Molecular Biology and Biotechnology, Faculty of Biology, Adam Mickiewicz University, 61-614, Poznań, Poland

**Keywords:** Transcription termination, pre-mRNA 3’ end processing, cancer progression, alternative polyadenylation

## Abstract

Two key factors required for pre-mRNA 3’ end cleavage and polyadenylation, CPSF73 and PCF11, exhibit oncogenic properties: their elevated expression is associated with poor cancer patient prognosis. However, both proteins are also transcription termination factors, and it is unclear which of the processes they promote might contribute to carcinogenesis. Here, we employ a cellular model of colorectal cancer (CRC) progression and find that cells from primary tumor are addicted to high levels of CPSF73 and PCF11, while metastatic cells become less sensitive to their levels. We find no association between alternative polyadenylation (APA) and cell dependence on CPSF73 and PCF11, and no impact of their downregulation on transcription-replication collisions. Instead, we uncover an uncoupling of changes in 3’ cleavage and termination during CRC progression: primary tumor-derived cells display a global shift to more proximal termination, yet a tendency for distal APA, compared to normal cells. Metastatic cells display partial reversion toward termination patterns observed in normal cells, and opposite tendency favoring proximal APA. This prompts us to measure the distance between the site of 3’ cleavage and transcription termination for active protein-coding genes and find it almost halved in cells from primary tumor compared to normal cells. Interestingly, this distance becomes critically short for oncogenes. Closer proximity of 3’ cleavage to termination correlates with higher gene expression, both across genes within a cell line and when distance and expression change in parallel. This uncovers a new relationship of transcription termination with gene expression regulation.

**SIGNIFICANCE STATEMENT:** At the 3’ end of genes two processes occur, which are often treated as one: pre-mRNA 3’ processing and transcription termination. In fact, they are promoted by a common set of protein factors, with oncogenic properties. Here, we employ a cellular model of colorectal cancer (CRC) progression, to study changes in both pre-mRNA 3’ processing and transcription termination. Unexpectedly, we uncover that those changes occur in opposite directions: mRNA tends to be lengthened, while termination is accelerated in cells from primary tumors. We show that the cleavage-termination distance is strongly reduced in these cells compared to normal and metastatic cells, and further determine that cleavage-termination proximity correlates with enhanced gene expression. This uncovers a new layer of gene expression regulation.

## INTRODUCTION

The annotated gene end is defined by the site of its pre-mRNA 3’ end processing. It is catalyzed by the cleavage and polyadenylation (CPA) complex and can occur at different alternative polyadenylation sites (PAS) within a single gene. The levels of CPA factors are frequently deregulated in various cancers. Many CPA factors, such as CPSF1, CPSF3, CPSF4, and CSTF2 are upregulated in different cancer types ^1–5^, while CFIm25 is downregulated in liver, breast, and cervical cancers^6–8^. In line with the changes in CPA factor levels, a global 3’ untranslated region (UTR) shortening and increased usage of intronic PAS are often observed in cancer cells^1,9–11^ However, in all cancer types tested, 3’ UTR changes are not unidirectional, and particularly abundant 3’ UTR lengthening events have been reported in colorectal and breast cancers^12,13^. Altogether, these findings highlight the complexity of cleavage and polyadenylation dysregulation in cancer.

The second crucial event at gene ends is RNA polymerase II (RNAPII) transcription termination. Termination occurs when RNAPII stops RNA synthesis and dissociates from the DNA template. It is necessary to separate adjacent RNAPII transcription units, both in tandem and convergent orientation, for co-regulating the expression of multiple genes^14–16^. Indeed, defective termination results in transcriptional interference on neighboring genes, which can lead to severe reduction in mRNA levels. Collided RNAPII are often removed from the DNA template through ubiquitylation-directed proteolysis^17^. Thus, it is not surprising that the production of readthrough transcripts in healthy human tissues is restricted in genomic regions with high gene activity^18^. Transcription termination of RNAPII-transcribed units is triggered by a whole plethora of complexes and protein factors. Some species of non-coding transcripts are terminated by Integrator and microprocessor complexes^19–22^. The Restrictor limits the expression of promoter upstream transcripts (PROMPTs) and early transcription^23–25^. Additionally, DNA-guided termination on T-tracts can function in RNAPII termination, as recently shown in yeast and humans^26,27^. This diversity of (partially redundant) mechanisms and complexes involved in RNAPII transcription termination underscores both the critical importance and the complexity of the process.

Transcription termination of protein-coding genes is mainly triggered by CPA factors and the XRN2 5’-3’ exonuclease^28,29^. In humans, this usually occurs several thousand nucleotides downstream of PAS^30–32^. Mechanistically, there are two key aspects of transcription termination on protein-coding genes, unified in a single model^33–36^. First, transcription of the polyadenylation signal induces an allosteric change in RNAPII and makes it competent for termination ^37^. Second, pre-mRNA 3’ end cleavage exposes the 5’ end of nascent RNA, allowing XRN2 to degrade it and help to displace RNAPII ^38–41^. Transcription termination is facilitated and preceded by RNAPII pausing, and terminally paused RNAPII can be detected by a specific phospho-mark on the C-terminal domain (CTD) of RNAPII’s largest subunit (RNAPII-CTD-T4ph)^30^. Two components of the CPA complex, CPSF73 and PCF11, are particularly important for transcription termination of protein-coding genes. CPSF73 is the endoribonuclease that cleaves the 3’ end of pre-mRNA^42^. Notably, rapid depletion of CPSF73 leads to extensive transcription readthrough, a hallmark of impaired termination^36^. PCF11 is essential for pre-mRNA 3’ cleavage both in yeast and humans^43–45^. It is also the only CPA factor shown to directly trigger transcription termination by binding to the RNAPII CTD and slowing it down ^32,46–51^. Besides stimulating normal termination on protein-coding gene ends globally, PCF11 also triggers premature termination within the introns of specific genes^32,52,53^.

Given their biological importance, it is not surprising that the levels and activity of PCF11 and CPSF73 are critical in cancer biology, and their manipulation has emerged as a promising therapeutic strategy^54^. Low PCF11 levels are associated with higher survival rates in neuroblastoma patients and spontaneous tumor regression^55^, as well as a better prognosis in hematologic malignancies^56^. Additionally, single nucleotide variants in the promoter of *PCF11* are recurrent in different cancer types ^57^. Similarly, pancreatic tumor patients with high *CPSF73* expression had significantly worse overall survival than patients with low *CPSF73* expression^5^. These findings justify the targeting of CPSF73 and PCF11 in cancer therapies. For instance, JTE-607 is a direct chemical inhibitor of CPSF73 activity, which was initially believed to inhibit cytokine production in human cells^58–60^. In a leukemia model, JTE-607 administration significantly prolonged mice survival^61,62^. The compound has also been reported to attenuate glioblastoma growth *in vivo* as well as breast and pancreatic cancer cell proliferation *in vitro*^5,63,64^. While CPA and termination factors CPSF73 and PCF11 emerge as novel targetable vulnerabilities for human cancers, it is unclear whether their roles in pre-mRNA 3’ processing or transcription termination are crucial for oncogenesis. Furthermore, the mechanisms linking their increased expression with poor patient prognosis remain obscure.

Colorectal cancer (CRC) is the third most diagnosed malignant tumor and the second leading cause of cancer deaths^65,66^. It is an etiologically heterogeneous disease that develops through three distinctive pathways: adenoma-carcinoma sequence, serrated pathway, and inflammatory pathway^67^. Although several critical genes have been reported to contribute to CRC development^68^, approximately 60–65% of CRC cases arise sporadically^67^. Due to CRC’s high prevalence, accompanied by poor tumor-selective drug delivery and low treatment efficacy in patients, there is a critical need to develop new therapies for CRCs^69^.

Here we employ a cell-line based model of colorectal cancer progression that exhibits elevated expression of CPA and termination factors, consistent with patient biopsies. We first show that the clonogenic potential of CRC cells critically depends on PCF11 and CPSF73, with cells from primary tumor displaying particularly high addiction to those factors. We then demonstrate that delaying transcription termination, either by genetic manipulation of termination factors or by chemical inhibition, does not increase the frequency of transcription-replication collisions, contrary to previous assumptions. Further on, we find no evidence supporting a role for alternative polyadenylation in CRC progression. Instead, by systematically characterizing the genomic landscape of RNAPII transcription termination and pre-mRNA 3’ end cleavage site usage in our model, we uncover an unexpected uncoupling between changes in transcription termination and in pre-mRNA 3’ end processing during CRC progression. To quantify this uncoupling, we introduce a new measure – distance between the cleavage site and the location in which RNAPII enters terminal pausing. We find that CRC cell sensitivity to CPA/termination factor depletion is associated with a shortened distance between pre-mRNA 3’ cleavage and transcription termination - particularly short in the case of oncogenes. Finally, we show that shortening of the cleavage-termination distance (i.e. increased proximity of cleavage and termination) on a gene is associated with its enhanced 3’ end formation and increased gene expression.

## RESULTS

### Primary and metastatic tumor cells show different sensitivities to RNA 3’ cleavage and transcription termination abrogation

To study the role of CPSF73 and PCF11 in oncogenesis, we employed a cellular model of colorectal cancer (CRC) progression: 1CT, non-transformed human colon epithelial cells^70^; HCT116, a widely used cell line derived from a primary tumor of the colon^71^; and two cell lines derived from the same patient: SW480 (from the primary tumor of the colon) and SW620 (derived one year later from metastasis to the lymph nodes)^72^ (Fig. 1A). This model recapitulates changes in the levels of the main players in pre-mRNA 3’ end processing and transcription termination of protein-coding genes: PCF11, CPSF73, and XRN2, as observed in patients. Proteomic data from biopsies of healthy colon and colorectal carcinoma^73^ indicated that the levels of all analyzed cleavage and polyadenylation (CPA) and termination factors, with exception of CLP1, increased in biopsies from colon cancer patients compared to normal colon tissue. This increase was observed in both absolute values as well as normalized to RNAPII (Fig. 1B, Fig. S1A). In line with the data obtained from patient samples, western blot analysis in our cellular model revealed a gradual increase of PCF11 protein levels concomitant with colorectal cancer progression (Fig. 1C, Fig. S1B). The levels of both CPSF73 and XRN2 were also elevated in CRC cells compared to 1CT cells but did not increase from primary to metastatic tumor cells (Fig. 1C, Fig. S1B). We conclude that protein levels of CPA and transcription termination factors are generally increased in CRC cells compared to normal colonic epithelial cells – both in our cellular model and in patient samples.

**Figure 1.**
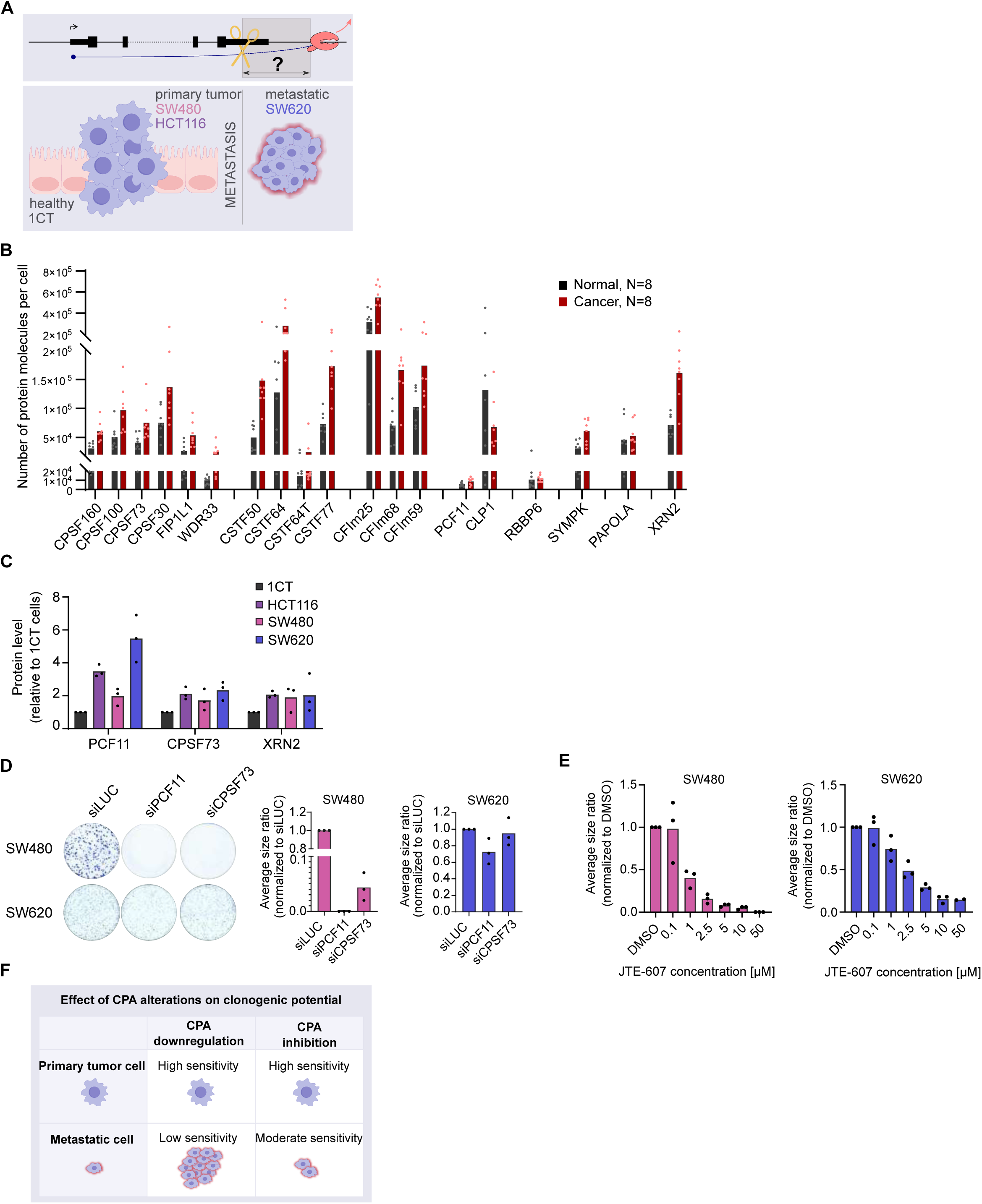
Primary CRC cells are more sensitive than metastatic cells to 3’ pre-mRNA cleavage and transcription termination abrogation. (A) Design of this study. The interplay between pre-mRNA 3’ cleavage and transcription termination was studied in a cell culture model of CRC progression. 1CT, non-transformed human colon epithelial cells; HCT116 and SW480, cells derived from colon primary tumor; SW620, cells isolated from metastasis in the SW480 donor patient. (B) Bar graph showing the number of protein molecules per cell of CPAand termination factors (data from patient biopsies^73^). (C) Bar graph showing the quantification of Western blots for PCF11, CPSF73, and XRN2 in the indicated cell lines. (D-E) Colony-formation assays in SW480 and SW620 cells. Cells were treated with siPCF11 or siCPSF73 (D), or with CPSF73 inhibitor JTE-607 (E). Here and in all other figures, bar graphs represent the average, while dots indicate individual biological replicates. (A) Schematic representation of the sensitivity of CRC cells to PCF11 or CPSF73 manipulation.

We then checked if and how the cancerous properties of primary tumor and metastatic CRC cells are affected by the loss of CPA and termination factor levels or activity. To address this, we used SW480 and SW620 cell lines, as they originate from the same patient (Fig. 1A). Knock-down of CPSF73 resulted in a strong inhibition of colony formation in SW480 cells, and PCF11 knock-down blocked colony formation altogether (Fig. 1D, Fig. S1C). In contrast, the clonogenic potential of SW620 cells was unaltered by CPSF73 downregulation and only slightly decreased upon PCF11 knock-down (Fig. 1D, Fig. S1C). Inhibition of CPSF73 endo-nucleolytic activity by JTE-607 at high concentrations abolished the ability of SW480 single cells to form colonies and decreased this ability for SW620 cells (Fig. 1E, Fig. S1D). This differential sensitivity of cells derived from primary and metastatic tumors in response to JTE-607 was consistent across a wide range of inhibitor concentrations (Fig. 1E, Fig. S1D).

These experiments suggest that the clonogenic potential of primary tumor-derived CRC cells is highly sensitive to changes in CPA factor levels and is readily impaired by their manipulation. In contrast, cells derived from CRC metastasis show a higher tolerance to CPA factor alterations and maintain their ability to form colonies (Fig. 1F).

### Induction of transcriptional readthrough by alteration of CPA factor levels or activity in CRC cells does not affect the frequency of transcription-replication collision

We next sought to understand the molecular background underlying the different sensitivities of primary tumor and metastatic CRC cell lines to varying levels of CPA/termination factors. Delayed termination has been speculated to increase the probability of collisions between RNAPII and replication machinery^74^. In line with this possibility, impaired pre-mRNA 3’ end cleavage and transcription termination were shown to cause reduction of replication fork speed and excessive origin firing^75^. This could lead to higher frequency of transcription-replication collisions (TRCs) and cell death. Before testing the hypothesis that CPSF73 and/or PCF11 downregulation increases the number of TRCs, we first assessed the number of TRCs in normal, primary, and metastatic tumor-derived cells. For this, we used the proximity ligation assay (PLA), which measures contacts between the transcription and replication machineries^76^. PCNA and T4ph RNAPII were used as markers for replication and transcription termination, respectively (Fig. 2A). Intriguingly, we observed a gradual decrease in TRCs in our cancer progression model, with the highest frequency in normal cells and the lowest in metastatic cells (Fig. 2A, right). However, when we manipulated CPA/termination factor levels or RNA 3’ cleavage activity, we did not observe the increase in TRC frequency predicted by our hypothesis. This result was consistent across multiple conditions: knock-down of PCF11 and CPSF73 in SW620 cells (Fig. 2B, left side), CPSF73 endo-nucleolytic activity inhibition by JTE-607 in SW480 and SW620 (Fig. 2B, right side), or CPSF73 depletion by an IAA-induced degron system in HCT116 cells^36^ (Fig. 2C, Fig. S2A). The only manipulation that resulted in a mildly elevated frequency of TRCs was depletion of XRN2 in HCT116 cells for 4 hours (but not after 8h)^28^ (Fig. S2B). The percentage of replicating cells (i.e. stained with EdU) after PCF11 or CPSF73 KD in SW620 cells also remained constant (Fig. S2C). To rule out potential bias from using the T4ph RNAPII-specific antibody in this assay, we additionally measured TRCs associated with Serine 2 phosphorylated (S2ph) elongating RNAPII and PCNA, as previously described^76–78^. Consistent with the previous results, neither knock-down of PCF11 or CPSF73 in SW620 cells nor treatment with JTE-607 in SW480 and SW620 cells led to changes in S2ph-linked TRC frequencies (Fig. S2D). Finally, we combined extended PCF11 knock-down (72 hours) with high-concentration JTE-607 treatment (50 µM) in SW620 cells, which still did not alter TRC frequency (Fig. S2E).

**Figure 2.**
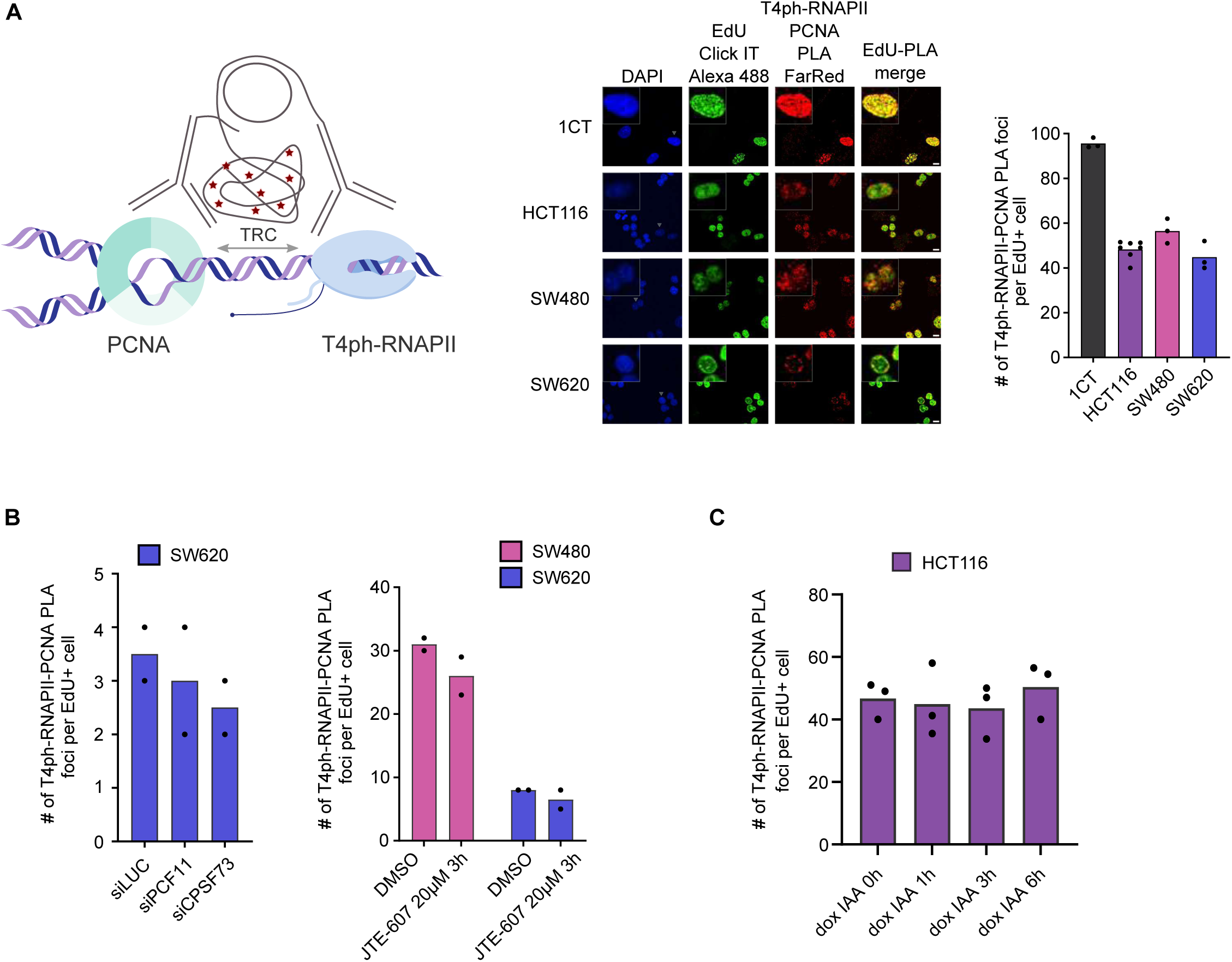
Frequency of transcription-replication collisions does not increase when transcription termination is disrupted. (A) Left: schematic representation of transcription-replication collisions (TRCs) quantification using proximity ligation assay (PLA). Right: representative pictures and TRC frequencies in the CRC model. Scale bar = 10 µm. (A-C) Bar graphs represent average values, dots individual biological replicates. (B) Frequency of TRCs in control conditions (siLUC, DMSO) and after knock-down of PCF11 (siPCF11) or CPSF73 (siCPSF73), or JTE-607 treatment in SW480 and SW620 cells. (C) Frequency of TRCs after CPSF73 depletion (dox IAA 0-6h) in HCT116 cells.

We conclude that PCF11/CPSF73 depletion and/or pre-mRNA 3’ cleavage inhibition, which were previously demonstrated to delay termination and induce transcriptional readthrough^32,79–81^, do not affect the number of transcription-replication collisions in CRC cells. Therefore, TRCs do not appear to underlie the sensitivity of CRC cells to PCF11/CPSF73 modulation in colony formation assays (Fig. 1D-F).

### Upregulation of CPA factor levels in CRC cells is associated with complex APA events instead of gradual mRNA shortening

A previous study investigating the effects of the JTE-607 inhibitor suggested that cells with higher CPA factor expression levels may undergo mRNA shortening, rendering them more sensitive to JTE-607^80^. To test this hypothesis in our model, we performed 3’ mRNA sequencing of nuclear RNA. Principal component analysis (PCA) revealed not only high reproducibility between biological replicates, but also a clear clustering and overlap of data from the two cell lines derived from primary tumors of different patients. This remarkable level of similarity in their PAS usage suggests a strong transcriptional memory of their tissue of origin (Fig. 3A). We then conducted alternative polyadenylation (APA) analysis to determine the direction of changes in 3’ end usage of CRC cells relative to 1CT (Fig. 3B, Fig. S3A; see also Methods). 3422 analyzed genes had a single PAS. Among genes undergoing APA, metastatic SW620 cells showed more proximal (22%, 2010 genes) than distal (15%, 1374 genes) APA events relative to 1CT. Strikingly, both primary tumor cell lines (HCT116 and SW480) displayed a clear opposite trend with a preference for distal (26%, 2393 genes and 34%, 3090 genes, respectively) relative to proximal (16%, 1455 genes and 19%, 1750 genes) APA events (Fig. 3B, Fig. S3A). Inspection of individual genes confirmed this analysis (Fig. 3C, Fig. S3B).

**Figure 3.**
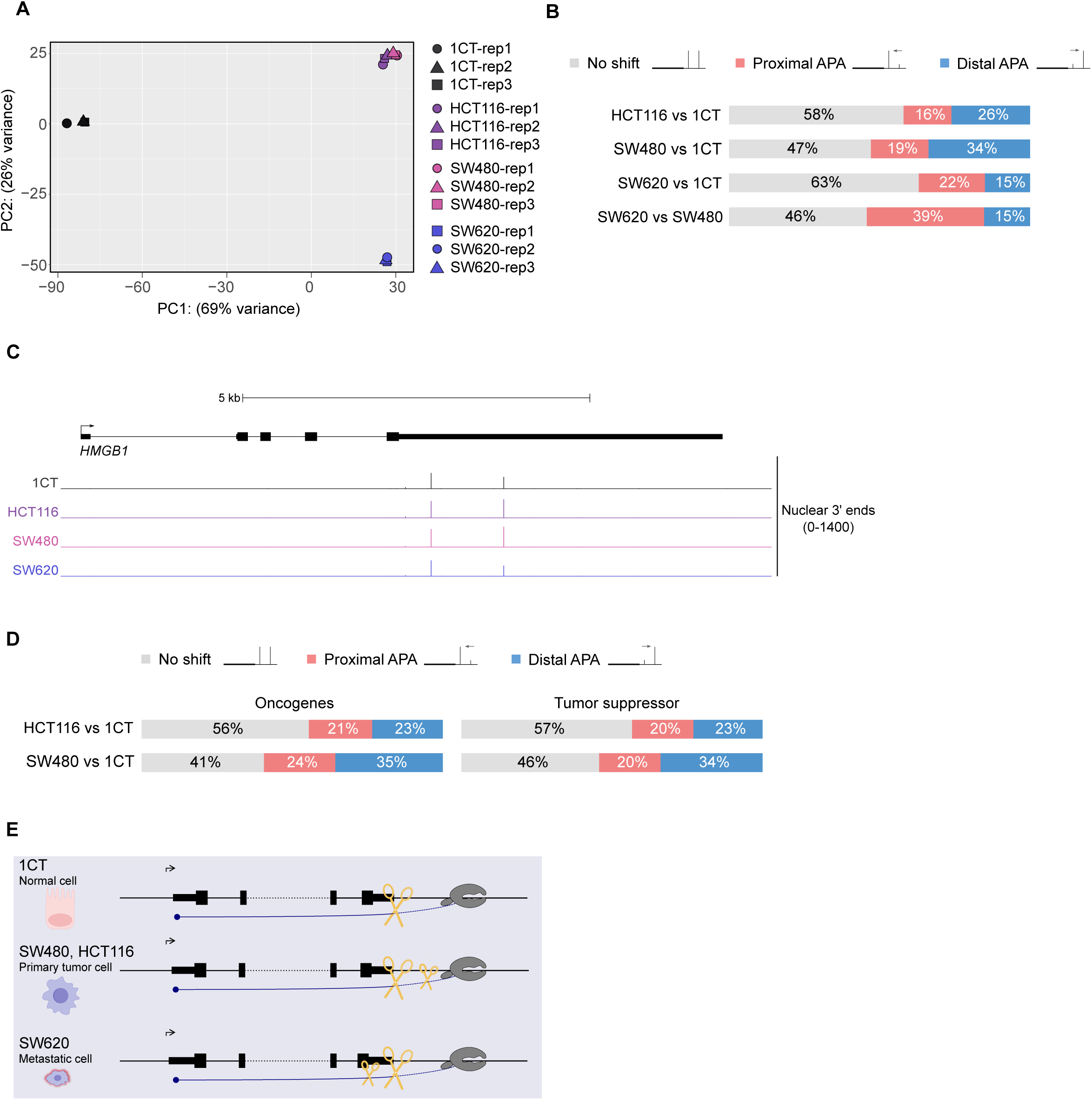
Changes in PAS usage in the CRC progression model. (A) Principal component analysis (PCA) plot of 3’ mRNA-seq biological replicates (N=3). (B) Bar graph representing the alternative polyadenylation (APA) analysis performed for separated active protein-coding genes (n=12469) in the CRC progression model. (C) Genomic profile of *HMGB1*, showing nuclear 3’ mRNA-seq signals. (D) Bar graph representing the APA analysis in the CRC progression model for separated oncogenes (n=533) and tumor suppressor genes (n=764) in primary tumor cells. (E) Schematic representation of PAS selection in the CRC progression model. The small scissors indicate APA direction.

We further checked whether genes involved in carcinogenesis show any tendency towards APA changes in the CRC model. We performed APA analysis for oncogenes and tumor suppressor genes. The results mirrored the trend observed for separated active protein-coding genes (see Methods), with no increase in APA incidence or directionality for cancer-related genes in CRC (Fig. 3B, D, Fig. S3A, C). Moreover, we did not observe a consistent correlation between APA and nuclear gene expression (Fig. S3D). Together, these findings show that upregulation of CPA factor levels is not associated with gradual mRNA shortening, but rather with complex, bi-directional APA events. They also suggest that in our model APA has no major global contribution to carcinogenesis or to the sensitivity of primary tumor cells to CPA factor alterations (Fig. 3E).

### Changes in PAS selection and transcription termination are uncoupled during CRC progression

Since CPSF73 and PCF11 are required not only for pre-mRNA 3’ cleavage but also for terminating transcription on protein-coding genes, we next investigated nascent transcription and transcription termination changes in our colorectal cancer model. We first globally examined nascent transcription profiles using mammalian native elongating transcript sequencing (mNET-seq) with an antibody recognizing all forms of RNAPII (total RNAPII)^82^. This revealed a proximal shift of nascent transcripts associated with total RNAPII (earlier termination) in PAS regions for cells derived from primary CRC, suggesting accelerated termination (Fig. 4A, C). To obtain a more specific and higher-resolution view of transcription termination, we next performed mNET-seq with an antibody against RNAPII T4ph - which detects RNAPII undergoing terminal pausing^30^. This method revealed that transcription termination downstream of the PAS (full-length termination) was indeed shifted proximally in CRC cells, most prominently in primary tumor cells, but also detectable in SW620 cells (Fig. 4B, C, F, Fig. S4D, E). Interestingly, full-length termination in metastatic cells shared characteristics of both normal and primary tumor cells (Fig. 4B, C, F, Fig. S4D, E), suggesting a partial reversal of termination patterns to those observed in normal cells. In addition to shifted gene-end termination, we detected more T4ph-marked RNAPII within gene bodies, suggesting an increase in the incidence of premature transcription termination in CRC cell lines (especially from primary tumors) compared to 1CT cells (Fig. 4B, D, Fig. S4A, B, E).

**Figure 4.**
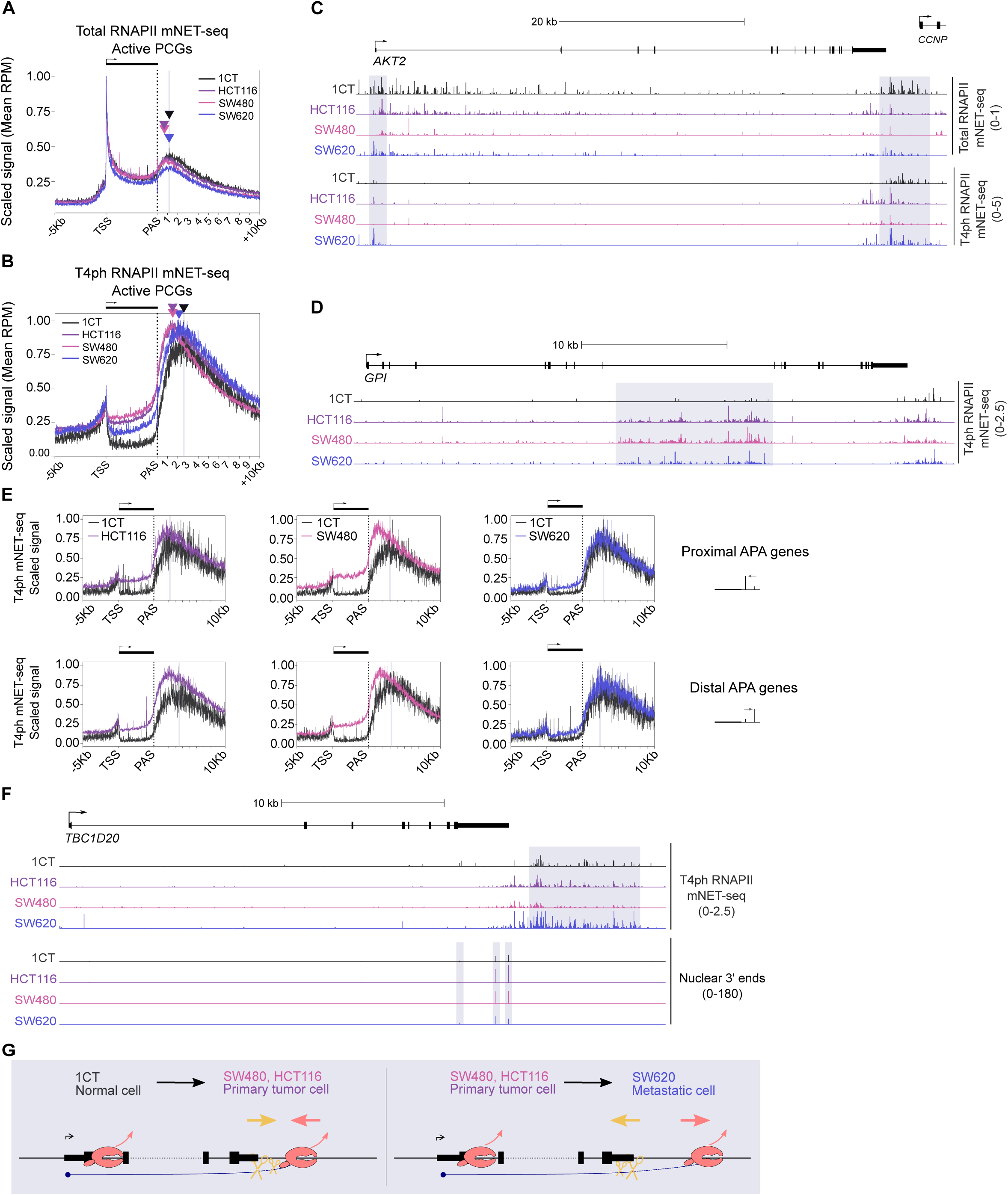
Changes in PAS selection and transcription termination are uncoupled during CRC progression. (A-B) Metagene profiles of Total RNAPII (A) and T4ph RNAPII (B) mNET-seq signal along active protein-coding genes (PCGs) (n=13577). The dotted line indicates the PAS position, and the grey highlight indicates the signal maximum downstream of PAS in 1CT cells. Triangles depict the signal maxima downstream of PAS. (C-D) Genomic profiles of *AKT2* (C) and *GPI* (D), showing Total and T4ph RNAPII mNET-seq signal. The grey highlight indicates RNAPII pausing in 1CT cells (C) or premature transcription termination region in CRC cells (D). (E) Metagene profile of T4ph-mNET-seq signal on genes undergoing proximal or distal APA in the CRC progression model. (F) Genomic profile of *TBC1D20*, showing T4ph-mNET-seq and nuclear 3’ mRNA-seq signals. The grey highlight indicates T4ph-RNAPII pausing in 1CT cells or PAS usage. (G) Schematic representation of uncoupling of changes is PAS usage (yellow) and RNAPII termination (red) in the CRC progression model. Left: transition from normal cell to primary tumor. Right: transition from primary to metastatic tumor. The arrows indicate the direction of change.

Taken together, CRC progression in our model is associated with increased premature termination and a proximal shift in full-length termination in cells from primary tumors compared to normal cells. Strikingly, metastatic tumor cells display a partial reversal of both premature and full-length termination, with a termination landscape in between the one observed in normal and primary tumor-derived cells.

The proximal shift in transcription termination in cell lines from primary tumor was surprising – because the direction of changes was opposite to the tendency toward distal APA in those cell lines (Fig. 3). To corroborate this and exclude the possibility that the proximal termination shift and distal APA occur mutually exclusively (i.e. on different subsets of genes), we checked transcription termination profiles on genes across all APA categories. This revealed accelerated termination (T4ph mNET-seq) in CRC cells compared to 1CT cells, regardless of presence/absence or even the direction of APA in the analyzed gene category (Fig. 4E, Fig. S4C). This accelerated transcription was visible as both increased premature termination in the gene body and proximally shifted full-length termination. Inspection of individual genes confirmed this analysis – earlier full-length termination yet distal PAS usage in cells derived from primary CRC can occur on the same gene (Fig. 4F, Fig. S4D). Admittedly, we also found examples where earlier transcription termination in primary CRC cells was accompanied by proximal PAS selection (Fig. S4E). Collectively, the above results suggest that the direction of APA and changes in transcription termination are globally uncoupled during colorectal cancer progression (Fig. 4G).

### Proximity of pre-mRNA 3’ end cleavage and transcription termination predicts upregulation of processed nuclear transcripts

The unexpected finding that changes in PAS selection and localization of RNAPII terminal pausing occur in the opposite direction during CRC progression prompted us to investigate their relationship more systematically. We quantified the distances between the genomic coordinate of the major PAS used for a given gene in each cell line, and the start coordinate of the region in which RNAPII enters terminal pausing mode (the onset of transcription termination) in each cell line. For simplicity, we refer to this measure as the cleavage-termination distance. Our analysis revealed that the median cleavage-termination distance in primary tumor-derived cells dropped sharply, to roughly half of the distance observed in normal cells, and partially recovered in CRC metastasis-derived cells (Fig. 5A). We also noticed that the cleavage-termination distance tends to be particularly short for oncogenes, as compared to both active protein-coding genes and tumor suppressor genes, for all cell lines of our model (Fig. 5B, Fig. S5A, B).

**Figure 5.**
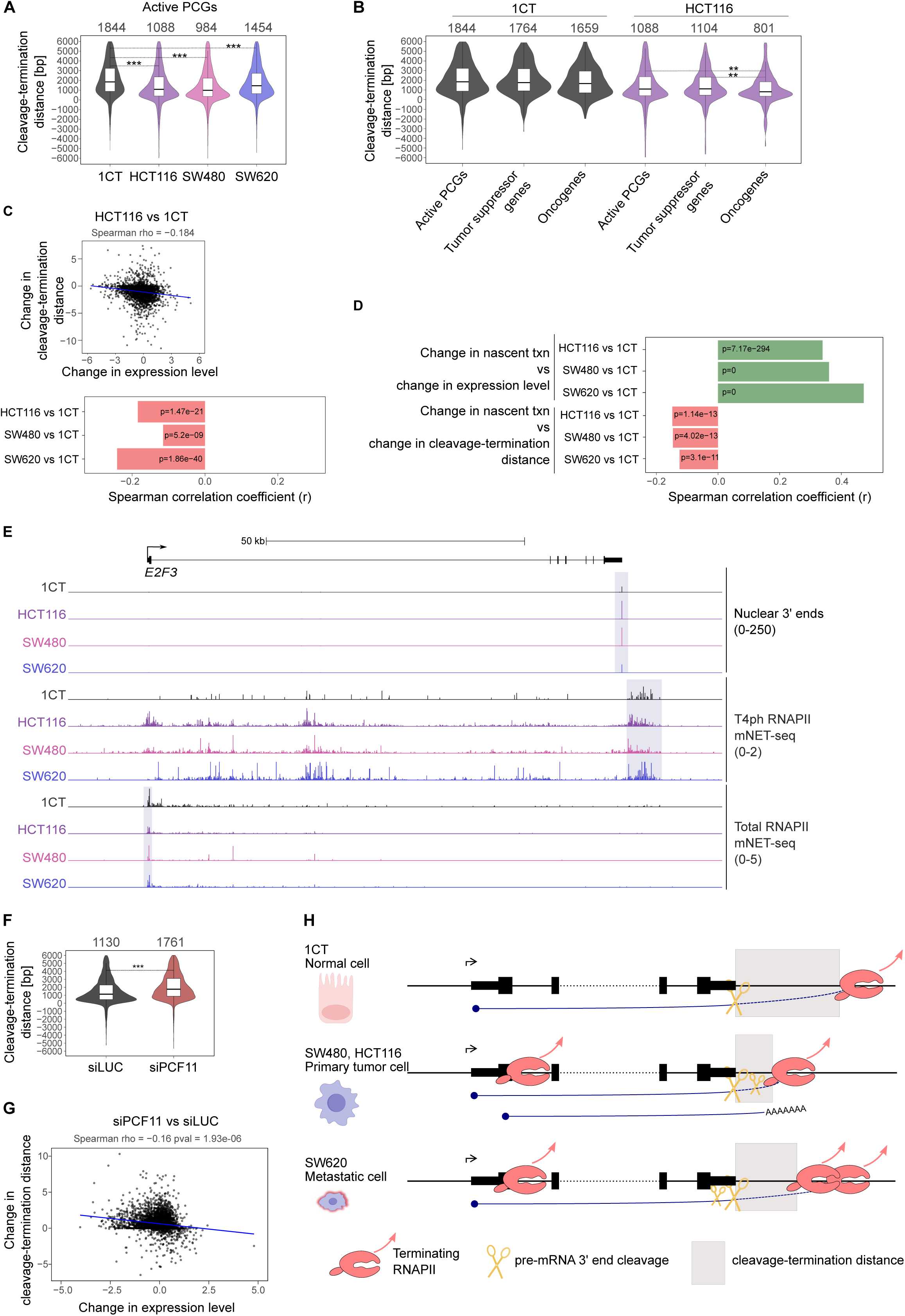
Closer proximity of PAS and transcription termination (onset of terminal pausing) is associated with higher gene expression. (A) Violin plot representation of the cleavage-termination distance for active PCGs (n=13577). (A-B) ***, Padj < 0.001; ** < 0.01; * <0.05. Non-indicated differences are not significant. The numbers above the plots represent the median. (B) Violin plot representation of the cleavage-termination distance for active PCGs, tumor suppressor genes, and oncogenes, in 1CT and HCT116 cells. (C) Pairwise correlations between changes in gene expression and the cleavage-termination distance in cancer cells compared to 1CT cells. The scatter plot comparing HCT116 and 1CT cells (upper panel) and the r and p-values for all the comparisons performed (lower panel). Positive values indicate increase in expression and cleavage-termination distance, negative values decrease. Each bar represents a pairwise comparison, its width indicates the Spearman correlation coefficient. Pink bars indicate negative correlation. (D) Pairwise correlations between changes in nascent transcription (txn), gene expression, and cleavage-termination distance in cancer cells compared to 1CT cells. Each bar represents a pairwise comparison, its width indicates the Spearman correlation coefficient. Pink bars indicate negative correlation, green bars indicate positive correlation. (E) Genomic profile of *E2F3*, showing nuclear 3’ mRNA-seq, T4ph-mNET-seq and Total-mNET-seq signals. The grey highlight indicates PAS usage and RNAPII pausing in 1CT cells. (F) Violin plot representation of the cleavage-termination distance for active PCGs in HeLa cells in control conditions (siLUC) or upon PCF11 knock-down (siPCF11) (n=11330). (G) Pairwise correlation between changes in gene expression and the cleavage-termination distance upon PCF11 knock-down in HeLa cells. Positive values indicate increase in expression and cleavage-termination distance, negative values decrease. (H) CRC progression is associated with more premature and proximal full-length transcription termination in primary tumor compared to normal cells. In metastatic cells, premature termination is still increased, but transcription termination downstream of PAS is partially reversed to the location in normal cells. Smaller scissors indicate APA direction: distal in primary tumor cells and proximal in metastatic cells. Primary tumor cells are characterized by shorter cleavage-termination distance that is associated with higher expression level.

We then wondered whether the cleavage-termination distance might influence gene expression. For the purpose of our study, we defined gene expression as the level of correctly processed transcripts measured by nuclear 3’mRNA sequencing (see Methods). All cell lines from our model displayed a slight but consistent tendency for lowered gene expression with increased cleavage-termination distance (Fig. S5C, darker violin plots). A small subset of active genes (7-11% per cell line) showed a negative cleavage-termination distance – terminal RNAPII pausing could be observed on those genes already proximally to the main cleavage site (Fig. S4D). We found negative cleavage-termination distance to be associated with lower gene expression (Fig. S5C, lighter violin plots), but did not study this small subset further. We then checked how changes in the cleavage-termination distance relate to changes in the expression of the same gene in our CRC model. For each of the CRC cell lines, an increase in cleavage-termination distance of a gene during progression from normal to cancer cell significantly anticorrelated with an increase of its expression (Fig. 5C, Fig. S5D). In simple terms: shortening of the cleavage-termination distance was associated with increased expression of the gene.

We next wanted to understand the mechanism behind increased expression of genes with shorter cleavage-termination distance. We hypothesized it could be driven by increased nascent transcription, more efficient RNA processing, or both. First, we checked how changes in nascent transcription (total RNAPII mNET-seq signal) of the gene impacted changes in gene expression (defined by nuclear 3’mRNA sequencing) in our CRC model, relative to 1CT. As expected, those two measures showed a highly significant (p=0) correlation (Fig. 5D, top panel). However, the correlation coefficient was below 0.5, indicating that transcriptional changes were not solely or dominantly responsible for the changes in levels of correctly processed nuclear transcripts. In line with this result, changes in nascent transcription of the gene significantly anticorrelated with changes in its cleavage-termination distance (Fig. 5D, bottom panel, Fig. S5E), although with a smaller correlation coefficient compared to changes in gene expression (Fig. 5C). This implied that the increase in gene expression associated with shorter cleavage-termination distance is partially due to increased nascent transcription, but also enhanced RNA processing. This interpretation is supported by the observation of individual oncogenes (*E2F3*, *CDK1* and *EZH2*) in CRC compared to normal cells. They showed shortened cleavage-termination distance and increased expression, yet without an increase in nascent transcription signal (Fig. 5E, S5F-G), implying increased efficiency of RNA processing.

CRC progression is associated with a whole plethora of events and hence gene expression or cleavage-termination distance changes during CRC progression cannot be attributed to any one factor. In order to evaluate the relationship between cleavage-termination distance and gene expression under more controlled conditions, we used a system which we previously characterized in depth. We triggered a downstream shift of the transcription termination window (termination delay) by depletion of PCF11 in the cervical cancer cell line HeLa ^32^. Reassuringly, both in control and knock-down conditions, we observed the same trend as in CRC: the shorter the cleavage-termination distance, the higher the expression level (Fig. S5H). Further on, PCF11 knock-down resulted in increased cleavage-termination distance (Fig. 5F). This lengthening of the cleavage-termination distance of a gene by PCF11 knock-down correlated significantly with its decreased expression level (Fig. 5G), and the correlation coefficient was similar to that observed in CRC progression (Fig. 5C). We corroborated this finding also on individual genes (Fig. S5I).

In summary, our data reveals that the distance between the site of 3’ pre-mRNA cleavage and RNAPII entering into terminal pausing mode is halved in cells from primary CRC, compared to normal cells, and partially restored in cells from metastasis. Shorter cleavage-termination distance is associated with elevated levels of the corresponding nuclear mRNA, partially due to elevated nascent transcription, and in part to enhanced RNA processing. We speculate that the observed spatial (and possibly also temporal) proximity of cleavage and termination could underline the high sensitivity of primary tumor-derived CRC cells to manipulations of CPA levels with respect to their colony formation ability (Fig. 1).

## DISCUSSION

CPA and termination factors PCF11 and CPSF73 emerged recently as having oncogenic properties, with their high levels being predictors of poor outcome in several cancer types^5,55,56^. However, the molecular mechanisms involved are unclear. Our study revealed that CRC cells derived from primary tumor or its metastasis differ in their sensitivity to alterations in PCF11 and CPSF73 levels and/or activity. While those alterations block colony formation from primary tumor cells, metastatic cells were less sensitive to CPA factor depletion (Fig. 1D) and RNA 3’ cleavage inhibition by JTE-607 (Fig. 1E, Fig. S1D). A previous study found a strong correlation of gene expression changes in response to JTE-607 treatment with PCF11 knock-down, pointing to a common transcriptomic signature^80^. The same study suggested that increased CPA factor levels lead to proximal PAS usage, which could sensitize cells to JTE-607 treatment. However, this hypothesis was based on the comparison of two unrelated cell lines (liver- and cervix-derived). Here, we analyzed four cell lines originating from the same tissue (colon) with a comprehensive set of genomic assays. Two of the analyzed cell lines – SW480 (from primary tumor) and SW620 (from its metastasis) – were even isolated from the same patient. The level of CPA and termination factors increased in our colorectal progression model (Fig. 1C, Fig. S1B). In contrast, changes in APA were not progressive: while we observed more distal APA events in cells from primary tumor compared to normal cells, metastatic cells were characterized by more proximal PAS events (Fig. 3A-C, Fig. S3A-B). Since cells from primary tumors, with tendency to distal APA, were more sensitive to JTE-607 and CPA downregulation, this contradicts the previous hypothesis^80^ and demonstrates that shortening of mRNA does not by itself sensitize cells to CPA inhibition. We additionally found that the clonogenic potential of CRC cells was more dependent on PCF11 than CPSF73, in both SW480 and SW620 cell lines (Fig. 1C, Fig. S1B). This is consistent with PCF11 being a limiting and regulatory factor within the CPA complex^32^. Coincidentally, it is also the only factor known to be able to directly trigger termination^48^.

Our results further refuted the hypothesis that the high sensitivity of primary tumor cells to PCF11 or CPSF73 knock-down results from increased frequency of transcription-replication collisions (TRCs) (Fig. 2, Fig. S2). We performed PCF11 KD, CPSF73 KD, acute CPSF73 depletion with degron, and CPSF73 inhibition, which all result in global transcription readthrough^28,32,79,80^ but did not increase TRC frequency (Fig. 2B-C, Fig. S2D-E). The only readthrough inducing manipulation which led to a modest increase in TRC frequency was 4h of acute XRN2 depletion (Fig. S2B). However, besides its role in termination, XRN2 resolves R-loops, which may block replication fork progression^83,84^. Two studies reported replication stress upon transcription termination crisis. Depletion of elongation factor SPT6 led to increased levels of elongating RNAPII in intergenic regions and accumulation of R-loops and DNA damage presented by gamma-H2AX foci^85^. Knock-down of CPA factor WDR33 resulted in Ser2-RNAPII readthrough, transcription-dependent reduced replication fork speed, and more replication initiation events^75^. However, the influence of RNAPII readthrough on TRC frequency was not measured before. Our data indicate that inducing transcription readthrough by altering CPA and transcription termination factor levels does not affect the number of TRCs in CRC cells. The unchanged TRC frequency in the presence of elongation-competent but termination-deficient RNAPII might be connected with the ability of RNAPII to redistribute pre-replication complexes to transcriptionally inactive regions before onset of replication^86,87^.

It is worth noticing that we observed more TRC foci in SW480 compared to SW620 cells, in all conditions tested (Fig. 2A-B, Fig. S2D). Since these cell lines were isolated from the same patient^72^, we speculate that in this patient CRC cells evolved during metastasis to deal with DNA damage by limiting TRC frequency. Changes in TRC frequency upon cancer progression have not been measured before, so it would be interesting to check in future studies if downregulation of TRCs occurs during oncogenesis also in other types of cancers.

One of the most surprising findings in this study is that changes in pre-mRNA 3’ processing and transcription termination in CRC progression were uncoupled and even occurred in the opposite direction (Fig. 4G). In comparison to normal cells, primary tumor cells had accelerated (proximal) transcription termination (Fig. 4A-C, G, Fig. S4D-E). At the same time, distal cleavage sites were used more often, even on the same gene (Fig. 3B-C, Fig. 4E-F, Fig. S3A-B). Then, during the progression of CRC from primary to metastatic tumor, the opposite was observed: termination was delayed, while APA changes tended to shift in the proximal direction.

The second key finding of our study is that changes in the cleavage-termination distance on a given gene correlated negatively with changes of its expression, measured by the level of its processed transcript in the nucleus (Fig. 5C, Fig. S5D). Thus, shortening of the cleavage-termination distance is associated with increased gene expression. While correlation does not imply causality, it is important to note that this relationship was true not only in the CRC progression model (Fig. 5C, Fig. S5D), but also upon a relatively short genetic manipulation (48h knock-down of PCF11), which primarily delays transcription termination (Fig. 5G, Fig. S5I). The increase in gene expression is partially due to an increase in nascent transcription (Fig. 5D, upper). We think that increased nascent transcription could be due to accelerated recycling of RNAPII for subsequent rounds of transcription. Changes that cannot be attributed to transcription, we reason are due to more efficient pre-mRNA 3’ processing. On some genes, enhanced processing efficiency likely plays a dominant role, as exemplified by the oncogenes shown in Fig. 5E, Fig. S5F-G, where no increase in nascent transcription is visible.

The median cleavage-termination distance in cells from primary tumor was shortened to about half in comparison with cells from healthy colon epithelium (Fig. 5A). We speculate that such shortened opportunity window for cleavage and transcription termination in primary tumor cells makes these cells more vulnerable to PCF11 or CPSF73 manipulation (Fig. 1D-F). Only half of genes containing multiple alternative PASs (and a third of all active genes), underwent any APA between 1CT and SW480 cells (Fig. 3B), yet termination shifted proximally in SW480 cells even if there was no APA (Fig. S4C). This implies that the high sensitivity of primary tumor cells to CPA alterations is more likely to be caused by earlier termination (a robust and unidirectional shift), rather than by APA (which happens only on a subset of genes, and in two opposite directions, with just a slight tendency towards distal sites).

Overall, our work demonstrates that the interplay between pre-mRNA 3’ end cleavage and transcription termination is more complex than previously appreciated. It also suggests that the contribution of APA to carcinogenesis might have been overestimated while instead the effect of altered transcription termination on cancer biology was underestimated. So far, a few studies have measured the extent of readthrough, which can be done with any nascent transcriptomics technique. Here, the use of termination-specific T4ph-mNET-seq^30^ has put us in the unique position of being able to check the distance between the PAS and the onset of RNAPII terminal pausing. Interestingly, this distance negatively correlates with the expression of the gene and becomes critically short for oncogenes in primary CRC cells, which opens a new area of investigation. It can help to uncover the biogenesis and function of mRNA 3’ ends and the role of transcription termination in cancer progression. We hope such molecular studies will aid in exploiting cancer’s vulnerability to CPA and termination factor levels and developing new oncological drugs that will be used precisely in the cancer types and stages where they can be most effective.

## METHODS

### Cell culture

1CT cells were cultured in DMEM High Glucose + Medium 199 (4:1), 2% FBS, EGF (25 µg / mL, Thermo Fisher Scientific, PHG6045), hydrocortisone (1 µg / mL, MERCK, H0888), insulin (10 µg / mL, MERCK, I3536), transferrin (2 µg / mL, MERCK, T2036), sodium selenite (5 nM, MERCK, S5261), gentamicin sulfate (50 µg / mL, MERCK, G1397) on Corning Primaria 100 mm dish at 37 °C, 7% CO_2_, 3% O_2_.

HCT116, SW480, SW620, and HeLa cells were cultured in DMEM High Glucose, 10% FBS, penicillin/streptomycin in 5% CO_2_ at 37 °C.

All cell lines were negative for mycoplasma.

### Colony formation assay

Cells were split into 6 well dishes, 1500 cells per well, growing in 10% FBS media, 2mL. For analysis of knock-down of PCF11 or CPSF73, cells were transfected with Lipofectamine™ RNAiMAX Transfection Reagent (Thermo Fisher Scientific, 13778150) and siRNAs (siLUC, siPCF11, siCPSF73, Dharmacon) at 15 nM concentration. For JTE-607 treatment, cells were split at 1500 cells per well and approximately 24h later were treated with either DMSO or various concentrations of JTE-607 (MERCK, SML2833). Then cell culture continued for 10 days. After that, the medium was discarded, and wells were rinsed once with PBS, stained with 500 µL of Crystal Violet (MERCK; C0775) solution per well, incubated for 2 minutes at room temperature. Then Crystal Violet was aspirated, and wells were washed 4 times with 2 mL of water. Once residual liquid dried, pictures were taken with Odyssey® M Imaging System. Pictures were analyzed with ImageJ. Three biological replicates were performed.

### Immunoblotting

40 µg of proteins were resolved by electrophoresis using 7.5% or 4-15% Mini-PROTEAN® TGX™ Precast Protein Gels (BioRad) and transferred onto nitrocellulose membranes. Blots were blocked with 4% non-fat milk in PBST (PBS, 0.1% Tween-20), incubated with the following primary antibodies: anti-PCF11 (Abcam, ab134391, 1:200; RRID: AB_2783786); anti-CPSF73 (Abcam, ab72295, 1:200; RRID:AB_1268249); anti-XRN2 (Thermo Fisher Scientific, A301-103A, 1:200; RRID:AB_2218876), and anti-rabbit IgG – IRDye 800CW (LI-COR, 926-32211; RRID:AB_621843) or anti-rabbit IgG – Peroxidase (MERCK, A0545; RRID:AB_257896). Three biological replicates were performed.

### mNET-seq

100 µL of sheep anti-mouse IgG Dynabeads (Thermo Fisher Scientific, 11202D) for total RNAPII (Hiroshi Kimura, CMA601; RRID: AB_2827956) or 50 µL protein G Dynabeads (Thermo Fisher Scientific, 10004D) for T4ph RNAPII (Acitve Motif, 61361; RRID:AB_2750848) per one sample were washed 2x with ice-cold NET2+Empigen buffer (50 mM Tris pH 7.4, 150 mM NaCl, 0.05 % NP-40, 1% Empigen BB), mixed with 5 µg of the antibody, incubated O/N in cold room, washed 2x with ice-cold NET2 buffer and put on ice.

1CT cells on fourteen Primaria 100 mm dishes or cancer cells on one 150 mm dish were washed with ice-cold PBS, harvested by scraping on ice, moved to a 10 mL NUNC tube, and centrifuged at 500 g for 5 min at 4 °C. The cell pellets were then resuspended in 4 mL of ice-cold HLB+N buffer (10 mM Tris pH 7.5, 10 mM NaCl, 2.5 mM MgCl_2_ and 0.5 % NP-40) and lysed for 6 min in ice. The cell lysates were underlaid with 1 mL ice-cold HLB+NS buffer (10 mM Tris pH 7.5, 10 mM NaCl, 2.5 mM MgCl_2_, 0.5 % NP-40, 10 % Sucrose) and centrifuged at 800 g for 5 min at 4 °C. The nuclear pellets were resuspended in 120 µL of ice-cold NUN1 buffer (20 mM Tris pH 7.9, 75 mM NaCl, 0.5 mM EDTA and 50 % Glycerol). The nuclear lysates were transferred to new tubes, mixed with 1.2 mL of ice-cold NUN2 (20 mM HEPES-KOH pH 7.6, 300 mM NaCl, 0.2 mM EDTA, 7.5 mM MgCl_2_, 1 % NP-40 and 1 M Urea, PhosSTOP (MERCK, 4906837001), protease inhibitors (MERCK, 04693159001)), vortexed at maximum speed and incubated on ice for 10 min (sample vortexed every 3-4 min). The chromatin was then centrifuged at 4000 g for 1 min at 4 °C, washed with 500 µL of ice-cold PBS, 100 µl of ice-cold 1x MNase buffer (NEB, M0247) and digested in 100 µL of MNase (40 units / µL) reaction buffer for 2-10 min at 37 °C in a thermomixer (1200 rpm). The digestion reaction was stopped by the addition of 12.5 µL of 0.2 M EGTA and centrifuged at 16000 g for 5 min at 4 °C. The 100 µL of supernatant was diluted 10x with 900 µL of the ice-cold NET2+Empigen buffer, and RNAPII conjugated beads were added and incubated for 1 h in cold room. The beads were then washed six times with 1 mL of the ice-cold NET2+Empigen buffer and 50 µL of the ice-cold PNKT buffer (1 x T4 PNK buffer, NEB, M0236S, and 0.1 % Tween 20). Washed beads were incubated in 50 µL PNK reaction mix (1 x PNKT, 1 mM ATP, 0.05 U/ml T4 PNK 3’ phosphatase minus (NEB, M0236S) in Thermomixer (1200 rpm) at 37 °C for 5 min, and then washed with the NET2+Empigen buffer. Next, RNA was isolated from beads by addition of 1ml of TRI Reagent™ Solution (ThermoFisher, AM9738), chloroform purification, and precipitated with isopropanol. The RNA was resuspended in 10 µL of Urea dye (7M Urea, 0.05 % Xylene cyanol, 0.05 % Bromophenol blue), denatured and resolved on 6% TBE-Urea gel (Thermo Fisher Scientific, EC68652BOX) at 180 V for 15-18 min. In order to size select 25-100 nt RNA fragments, a gel fragment was cut between BPB and XC dye markers. A 0.5 mL tube was prepared with 4-5 small holes made with 25G needle and placed in a 1.5 mL tube. Gel fragments were placed in the layered tube and broken down by centrifugation at 16000 g for 1 min. The RNA fragments were eluted from gel using the RNA elution buffer (1 M NaOAc and 1 mM EDTA) at 25 °C for 1 h in Thermomixer (1200 rpm). The slurry was put into SpinX column (Costar, Corning, 8160) with 2 glass filters (Whatman, WHA1823010) and the flow-through RNA was ethanol precipitated. mNET-seq libraries were prepared using NEBNext Small RNA Library Prep Set for Illumina (NEB, E7330L). 15-18 cycles of PCR were used to amplify the libraries. Before sequencing, the libraries were size-selected on a 6%TBE gel (Thermo Fisher Scientific, EC62652BOX) selecting only the 150-230 bp PCR product to exclude primer dimers. Gel elution was performed as described above. The libraries were sequenced on NextSeq500 using NextSeq High-Output Kit, 75 cycles (Illumina) or NovaSeq 6000. 2-3 biological replicates were performed for each cell line.

### 3’ mRNA seq

The nuclei isolation was performed as for mNETseq experiments. The nuclear pellets were washed with 1 ml of ice-cold PBS and suspended in 500 µL of TRI Reagent™ Solution (Thermo Fisher Scientific, AM9738) and RNA was purified and ethanol-precipitated according to manufacturer’s instructions. DNA was removed by incubation with 4U of TURBO Dnase (Thermo Fisher Scientific, AM2238) at 37°C for 30 minutes and another round of TRI Reagent™ Solution – based purification. The RNA integrity was monitored by Agilent Tapestation. The libraries were prepared using Lexogen QuantSeq 3’ mRNA-Seq Library Prep Kit REV according to manufacturer’s protocol. 500 ng of RNA was used as an input. 12-13 cycles of PCR were used to amplify the libraries. Libraries size was controlled by Agilent Tapestation. Indexed libraries were quantified, pooled for sequencing, and sequenced on Illumina NovaSeq 6000. Three biological replicates were performed.

### Transcription-replication collisions

All PLA reactions were performed as described in ^88^ with a few modifications. Briefly, cells were seeded, 30 000 per well, in 96-well culture plates (Ibidi, 89626), grown as described previously and treated the following day.

For the knock-down experiment, the cells were transfected according to reversed transfection protocol. Final concentration of siRNA in transfection reaction reached 15 nM and 5uL of lipofectamine RNAiMAX (Thermo Fisher Scientific, 13778150).

To inhibit CPSF73, cells were treated with JTE-607 20 µM or 50 µM for 3h or 7h, respectively.

For CPSF73 depletion, cells were pretreated with doxycycline (1 µg/ml, MERCK, D9891) for 18h and next treated with 1 µM IAA (MERCK, SML3574; multiple timepoints). For XRN2 depletion, cells were treated with 1 µM IAA (multiple timepoints).

Before fixation, media was supplemented with 10µM EdU for 30 min for S phase identification. Immediately after, cells were washed 2x with PBS and pre-extracted for 2 min at RT using PBS + 0.5% Triton X-100. 1CT cells were pre-extracted for 10 min. After pre-extraction, the cells were fixed with 4% PFA for 15 min at RT, washed 3x with PBS, permeabilized for 1h with PBS + 0.5% Triton X-100, washed 3x with PBS, and then blocked with a PBS + 5 % BSA solution for overnight at 4 °C.

For S phase identification, EdU was labelled with AlexaFluor 488 Azide using a Click-it chemistry reaction as described in ^88^. Then cells were washed 2x with PBS and blocked for 1 h in PBS + 5 % BSA at RT. PLA reactions were performed following the manufacturer’s protocol (Merck) using the Duolink In Situ Detection Reagents FarRed kit (MERCK, DUO92013) and a mouse anti-PCNA antibody (Santa Cruz Biotechnology, sc-56,1:200; RRID:AB_628110) in combination with either a rabbit anti-(phospho-Ser2) RNA polymerase II antibody (Abcam, ab193468, 1:500; RRID:AB_2905557) or a rabbit anti-(phospho-Thr4) RNA polymerase II antibody (Novus biologicals, NBP1-49546, 1:100; RRID:AB_10011602) as primary antibodies. Lastly, a DAPI staining was performed by incubating the cells in a PBS + 5 µg/mL DAPI for 1 h at RT. Cells were then washed 3x with PBS and 400µL of PBS was left in the wells.

134 Z-stack images (11 µm, 1 image per µm) per well were acquired using Nikon T2 inverted microscope with an Andor Dragonfly spinning disk, combined with a Plan Apochromat 40X air objective with a numerical aperture of 0.95, a 2.0X magnification lens, and an iXon Life 888 EMCCD camera. Images were analyzed using ImageJ and an automated image analysis pipeline as described in ^88^. Briefly, the Z-plane being the most centered on nuclei is selected from the stack by identifying the highest DAPI variance plane. Then, the camera background signal is subtracted, nuclei are segmented using the DAPI signal, the Stardist Plugin, and a trained segmentation model ^89,90^, and finally fluorescence signals are measured, and PLA foci are counted using the ImageJ Maxima Finder tool. 2-6 biological replicates were performed for each experiment.

### Data Analysis

#### Genomic annotation and analyzed gene sets

Hg38/GRCh38 was used as a reference genome for mNET, and 3’mRNA-Seq analyses with basic gene annotation based on GENCODE release 43^91^. For most analyses, we selected protein-coding genes (PCGs) with at least 3 reads in the 1CT 3’mRNA data (see section “3’ mRNA-Seq”) and named them “active PCGs" (n = 13577). Using these genes, we customized annotations by assigning the TSS of the gene based on GENCODE coordinates, while the end coordinate corresponded to the experimentally detected PAS in 1CT 3’mRNA data. For genes with multiple PAS, we chose the end coordinate as the major PAS i.e., the PAS with the highest read count. This annotation was further used to extract subsets and generate metagene profiles.

For APA analysis, genes from the active PCGs gene set that did not overlap with another annotated gene on the same strand, and had a 3′ end isolated by at least 6 kb from the downstream annotated gene on the same strand were selected. This resulted in 12649 PCGs, referred to as "separated active PCGs”.

Carcinogenesis-related genes, including general oncogenes and tumor suppressor genes, were obtained from the oncogene (ONGene) (https://bioinfo-minzhao.org/ongene/)^92^ and tumor suppressor gene (TSGene) databases (https://bioinfo.uth.edu/TSGene/)^93,94^, respectively, yielding 679 oncogenes and 986 tumor suppressor PCGs. Further filtering for APA analysis within the active separated PCGs set identified 533 onco- and 764 tumor suppressor PCGs (Fig. 3D, Fig. S3C, S5A-B).

#### 3’ mRNA-Seq

3’ mRNA data sequenced on NovaSeq 6000 were converted to FastQ files using a illumina software bcl2fastq (v2.20.0.422). The files were then processed based on the specifications provided by Lexogen (https://github.com/Lexogen-Tools/quantseqpool_analysis). Aligned BAM files were then separated based on strand using SAMtools (v1.10). Further downstream analysis (below) was adapted from the previously published workflow for 3’ mRNA-Seq analysis^32^.

For PAS calls and APA analysis (below), we have adapted previously published workflows ^95–97^.

##### Filtering internal primming events

To minimize false positives caused by internal priming of the QuantSeq assay on A/T-rich genomic regions, we generated a basic genomic mask. This mask flags the areas in the genome with sequences containing six or more consecutive A’s (forward strand) or T’s (reverse strand), as well as any 10-nucleotide windows with more than six A’s or T’s. Next, to ensure that genuine polyadenylation sites (PAS) falling in A/T-rich regions were not included in the mask, we refined the mask by unmasking a 20-nucleotide window around/centered at each gene’s 3’ end (GENCODE v43) as well as experimentally validated PAS sites from previous studies^95^. Aligned 3’ mRNA-seq reads that overlap with the strand-specific refined masks were filtered out, and the remaining reads from the most distal nucleotide were retained for downstream analysis.

##### Polyadenylation site (PAS) Detection and Quantification

To identify polyadenylation sites (PAS) accurately, we used the filtered 3’ mRNA-seq reads (from the step described above) to create strand-specific, depth-normalized density profiles across the genome. These profiles were generated by summing up coverage signals from all 3’ mRNA-seq samples separately for each strand. Based on these profiles, PAS events were detected as follows: First, all signals within a 30-nucleotide window were summed up, and the windows with values above 30 were considered further. Within each window, the peak with the strongest signal was identified, and a new 30 nucleotide region was created around it. This step was repeated to eliminate overlapping intervals, yielding a set of reliable polyadenylation sites that were used to retrieve polyadenylation sites for each sample and to analyze the usage of alternative polyadenylation sites (APA).

##### Alternative PolyAdenylation (APA) analysis

For APA analysis, DEXSeq ^98,99^ was employed along with the previously published workflow for APA analysis^32^. Protein-coding genes that were considered active in 1CT 3’mRNA data, non-overlapping with any other gene on the same strand (n=12649; see "genomic annotation and analyzed gene sets") were subject to differential PAS usage quantification with DEXSeq. Gene coordinates were extended by 6kb downstream of the 3’ end (1CT major PAS) to allow for the detection of distal APA beyond gene ends (1CT major PAS). DEXSeq returns the fold change (log2) of the experimental sample over the control and adjusted p-value for each PAS. Genes where no PAS reaches significance (padj> = 0.05) were categorized as APA no shift. To assign the direction of APA shift in genes with multiple PAS, two most statistically differentially used PAS were retrieved (padj< 0.05). If the ratio of the distal to proximal site usage was higher in the experimental than in the control cells, the shift in the experimental sample was classified as distal, in the opposite situation as proximal.

##### Differential Expression analysis

Gene expression was quantified by summing nuclear 3′ mRNA-seq reads mapping to polyadenylation sites (PAS) within each gene’s 3′UTR, extended 6 kb downstream (to capture potential readthrough transcripts). These counts were then subjected for differential expression analysis with DESeq2^100^. DESeq2 outputs normalized counts for both experimental and control samples, along with the fold change (log2) of the experimental sample over control and an adjusted p-value for each gene. These values were further used for correlation analysis.

#### mNET-Seq

For mNET-Seq analysis, both T4ph and Total paired-end reads in fastq files were first subjected to quality control (FastQC v0.11.9), and then the adapter sequences were trimmed using TrimGalore (v0.6.6). Trimmed reads were aligned to the human genome using STAR^101^ (v2.7.6a), only allowing for one alignment to the reference. Strand-specific BAMs were generated using SAMtools (v1.10), and the replicates were merged into a single BAM using the same. To obtain single nucleotide resolution of strand-specific BAMs, the last transcribed nucleotide position from each aligned read was retrieved and RPM normalized using an in-house R script based on Bioconductor libraries.

#### Cleavage-termination distance analysis

To quantify cleavage-termination distances, we first identified termination windows - genomic regions enriched for T4ph-specific mNET-seq signal – using the MACS2 (v2.2.7.1) bdgbroadcall function on strand-specific bedGraph files with default parameters. These windows were then used to calculate the distance between each gene’s major PAS (gene end) and the start of the termination signal - cleavage-termination distance. We excluded the windows that began within the gene body and retained only those windows starting at the 3’UTR and within 6 kb of a gene’s major PAS. This resulted in two categories of distances 1) Negative – when the termination window begins upstream of a gene’s major PAS; 2) Positive – when the termination window begins downstream of the gene’s major PAS. For each cell line its corresponding major PAS was used and the analysis was performed for both active PCGs and carcinogenesis-related genes. Distances were then visualized in R using the ggplot2 (v3.4.4) function.

#### Metagene profiles

Metagene profiles were plotted on the active PCGs and different APA category gene sets. The profiles were generated using computeMatrix and plotProfile functions from the deepTools suite (v.3.5.1).

To scale the metagene profiles to 1, for each profile, a tab file consisting of numerical data underlying the plot was retrieved by specifying an option “--outFileNameData” while plotting using plotProfile. This tab file was used to extract the maximum value for each sample on the plot. Then, we used this maximum value to calculate a scaling factor = 1/max. The scaling factor of each sample was multiplied with its corresponding unscaled dataset (matrix). The profiles were then plotted again using the scaled dataset as an input for plotProfile and customized in R using ggplot2 (v3.4.4) function.

#### Correlation analysis

Correlation analysis was performed across gene expression, nascent expression, APA and cleavage-termination distance. Nascent transcription levels were quantified by extracting the total RNAPII signal from the transcription start site (TSS) to the 1CT major PAS for each gene using metagene profiles. For each gene, log2 relative values were computed for colorectal cancer (CRC) cells (HCT116, SW480, and SW620) relative to 1CT, and for HeLa siPCF11 cells relative to siLUC. These log2-relative values were then used to compute pairwise Spearman correlations among the colorectal model cells or HeLa cells.

Relative values are defined as below: gene expression was quantified as the log2 fold change of CRC vs 1CT (DESeq2), with negative values indicating downregulation and positive values as upregulation; APA was represented as the log2(distal/proximal) difference between CRC and 1CT (DEXSeq) (log2[CRC_distal/proximal] – log2[1CT_distal/proximal]), where negative values indicate a shift towards proximal PAS usage and positive towards distal PAS usage; cleavage-termination distance: log2 difference in distance between CRC and 1CT (log2[CRC_distance] – log2[1CT_distance]), with negative values representing shorter and positive as longer distances; and nascent transcription: log2 difference in total RNAPII signal between CRC and 1CT (log2[CRC_signal] – log2[1CT_signal]), where negative values indicate lower and positive as higher nascent transcription signal/levels.

## ACKNOWLEDGMENTS

We thank Nick Proudfoot, Joanna Kwiatkowska and Mariia Kapeliukha for critical reading of the manuscript. This work was funded by the European Union, Horizon Europe, ERC grant AlternativeEnds project number 101042642, National Science Centre, Poland, project number 2018/30/E/NZ1/00073, Polish National Agency for Academic Exchange project PPN/PPO/2018/1/00053, and European Molecular Biology Organization EMBO IG 4728-2020. M.R.G was supported by National Science Centre, Poland, project number 2018/31/B/NZ2/04004. S.H. was supported by the Helmholtz Association and the European Research Council (ERC Starting Grant 852798 ConflictResolution).

## AUTHOR CONTRIBUTIONS

Agata Stępień: Conceptualization; Data curation; Formal analysis; Investigation; Methodology; Visualization; Writing – original draft; Writing – review & editing.

Deepshika Pulimamidi: Data curation; Formal analysis; Investigation; Methodology; Visualization; Writing – original draft; Writing – review & editing.

Martyna Plens-Gałąska: Formal analysis; Investigation; Methodology; Visualization; Writing – review & editing.

Maxime Lalonde: Formal analysis; Investigation; Methodology; Visualization; Writing – review & editing.

Magda Kopczyńska: Investigation, Writing – review & editing. Hiroshi Kimura: Resources, Writing – review & editing.

Shazia Irshad: Resources, Writing – review & editing.

Michał Gdula: Methodology.

Stephan Hamperl: Investigation, Funding acquisition.

Kinga Kamieniarz-Gdula: Conceptualization, Data curation; Funding acquisition; Investigation; Methodology; Supervision; Visualization; Writing – original draft; Writing – review & editing

## DISCLOSURE AND COMPETING INTEREST STATEMENT

The authors declare no competing interests.

### Re-analysed data

In addition to the datasets generated in this study, we also re-analyzed previously deposited data. HeLa_PCF11 knock-down T4ph mNET-Seq and 3’mRNA-Seq are part of the super series GSE123105. The accession IDs for these samples are provided in the Supplementary Table1.

**Suppplementary_Table1:** List of sample accession IDs corresponding to the re-analysed data.

| Supplementary_Table1: List of sample accession IDs corresponding to the re-analysed data |  |  |  |
| --- | --- | --- | --- |
| Sample | Sample Accession Id | GEO Accession Id | Condition |
| T4ph_mNETseq_siLUC_rep1 | GSM3495849 | GSE123104 | Control |
| T4ph_mNETseq_siLUC_rep2 | GSM3495850 | GSE123104 | Control |
| T4ph_mNETseq_siLUC_rep3 | GSM3495851 | GSE123104 | Control |
| T4ph_mNETseq_siPCF11_rep1 | GSM3495852 | GSE123104 | KnockDown |
| T4ph_mNETseq_siPCF11_rep2 | GSM3495853 | GSE123104 | KnockDown |
| T4ph_mNETseq_siPCF11_rep3 | GSM3495854 | GSE123104 | KnockDown |
| 3'mRNAseq_siLUC_rep1 | GSM3536477 | GSE123105 | Control |
| 3'mRNAseq_siLUC_rep2 | GSM3536478 | GSE123105 | Control |
| 3'mRNAseq_siLUC_rep3 | GSM3536479 | GSE123105 | Control |
| 3'mRNAseq_siLUC_rep4 | GSM3536480 | GSE123105 | Control |
| 3'mRNAseq_siPCF11_rep1 | GSM3536481 | GSE123105 | KnockDown |
| 3'mRNAseq_siPCF11_rep2 | GSM3536482 | GSE123105 | KnockDown |
| 3'mRNAseq_siPCF11_rep3 | GSM3536483 | GSE123105 | KnockDown |
| 3'mRNAseq_siPCF11_rep4 | GSM3536484 | GSE123105 | KnockDown |

## Supplemental figures

**Figure S1.**
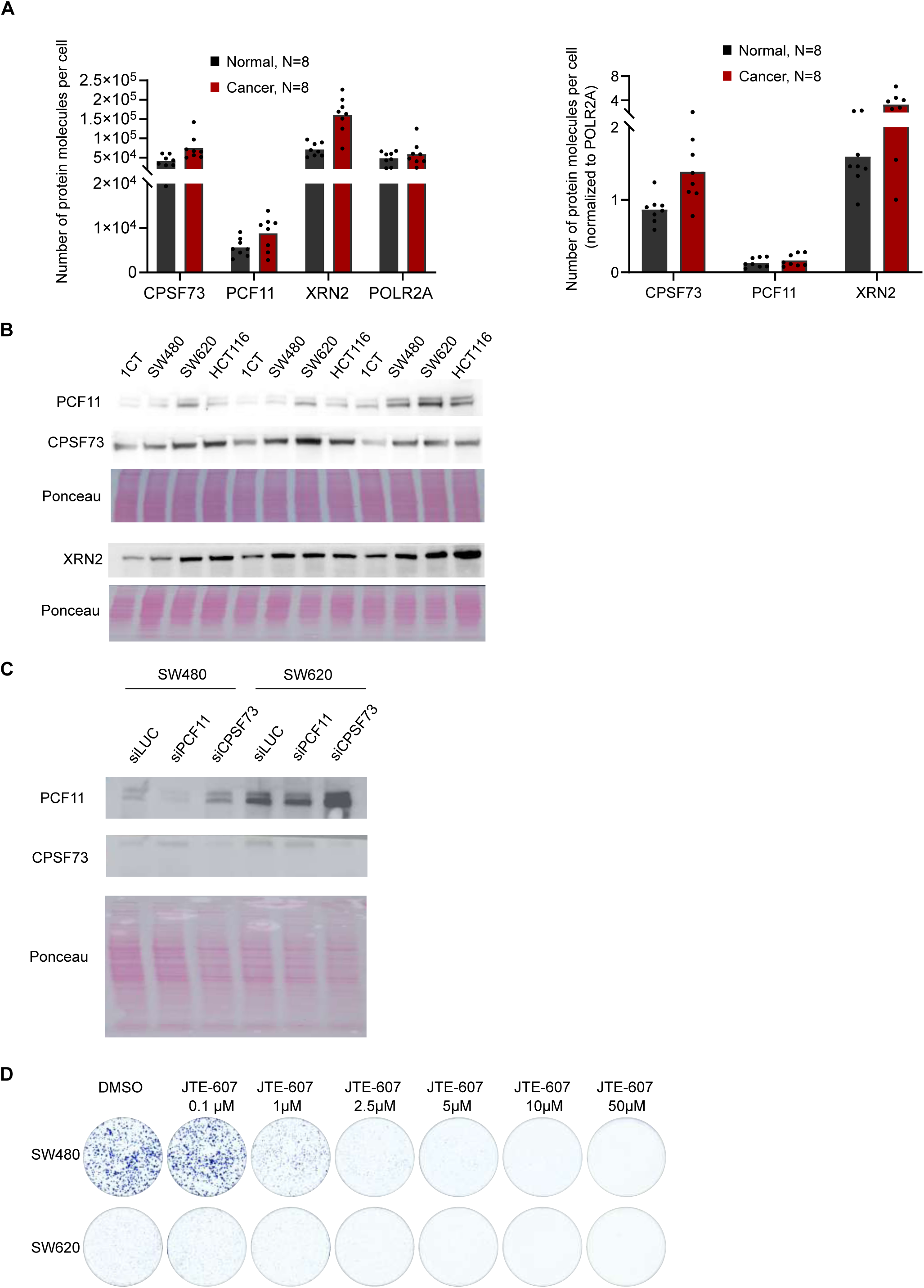
Protein levels of PCF11, CPSF73, and XRN2 in CRC and the effect of their manipulations on clonogenic potential of CRC cells. Related to Figure 1. (A) Bar graph showing the absolute (left panel) or normalized (to the largest subunit of RNAPII; right panel) number of protein molecules per cell based on global quantitative proteomics from patient biopsies^73^. Individual dots represent individual biological replicates. (B) Western blot showing the levels of PCF11, CPSF73, and XRN2 in the cell-culture based CRC progression model. (C) Western blot showing the levels of PCF11 and CPSF73 after PCF11 or CPSF73 knock-down (siPCF11 or siCPSF73) in SW480 and SW620 cells. (D) Representative examples of colony-formation assays in SW480 and SW620 cells treated with different concentrations of JTE-607, quantified in Figure 1E.

**Figure S2.**
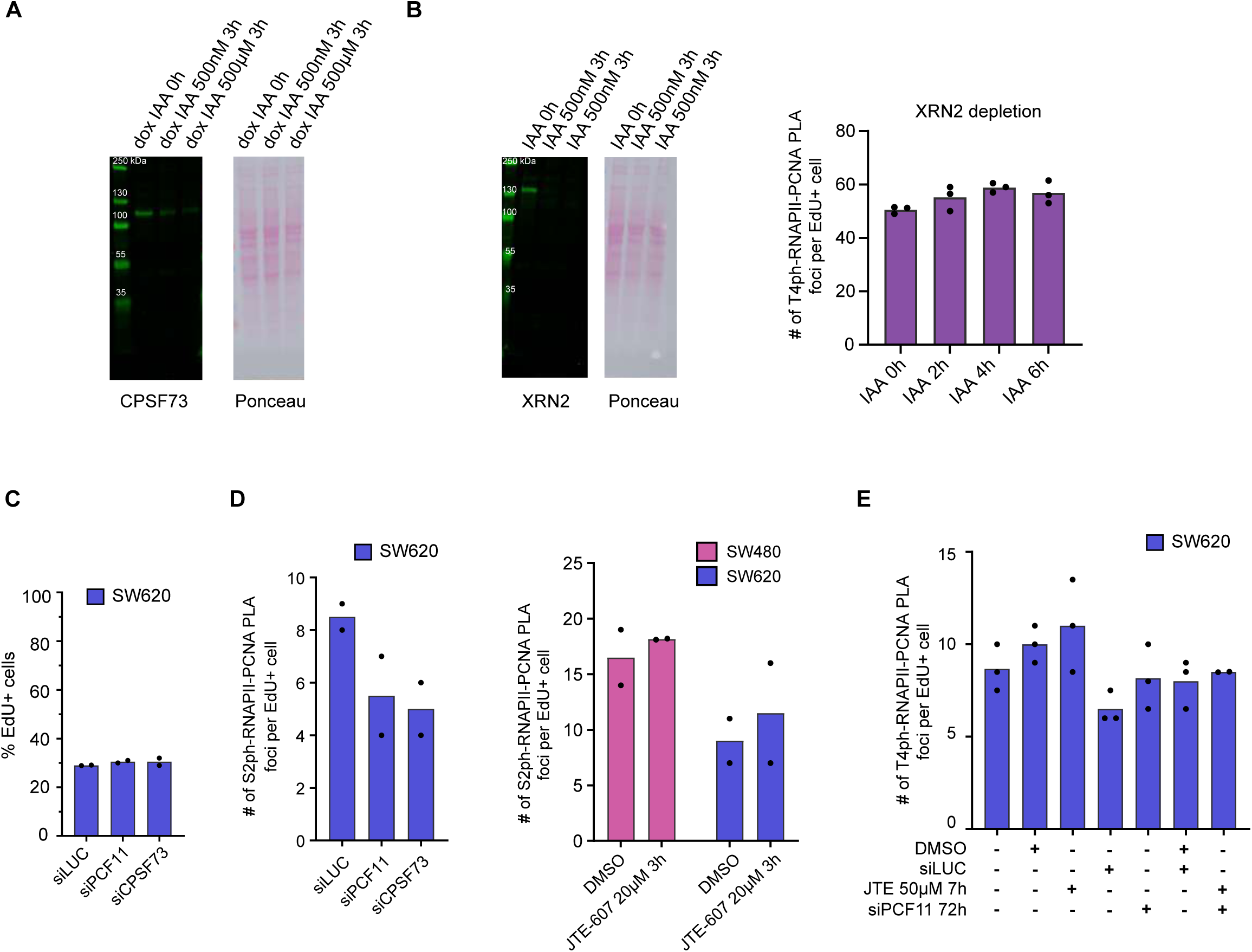
Frequency of transcription-replication collisions upon transcription termination factors alterations. Related to Figure 2. (A) Western blot showing the level of CPSF73 after acute CPSF73 depletion (dox IAA 3h) in HCT116 cells. (B) Western blot showing the level of XRN2 after acute XRN2 depletion (IAA 3h) and bar graph representing the number of transcription-replication collisions (TRCs) after XRN2 depletion (IAA 0-6h) in HCT116 cells. (B-E) Dots represent individual biological replicates. (C) Bar graph showing the % of EdU-positive cells in control (siLUC), PCF11 knock-down (siPCF11) or CPSF73 knock-down (siCPSF73) conditions in SW620 cells. (D) Bar graphs showing the number of TRCs in control (siLUC) or PCF11 (siPCF11) or CPSF73 (siCPSF73) knock-down conditions in SW620 cells (left) or after treatment of SW480 or SW620 cells with JTE-607 (right) obtained with the use of antibody against Ser2-phsphorylated RNAPII. (E) Bar graph showing the number of TRCs after prolonged treatment of SW620 cells with siPCF11 or JTE-607 or combined treatment.

**Figure S3.**
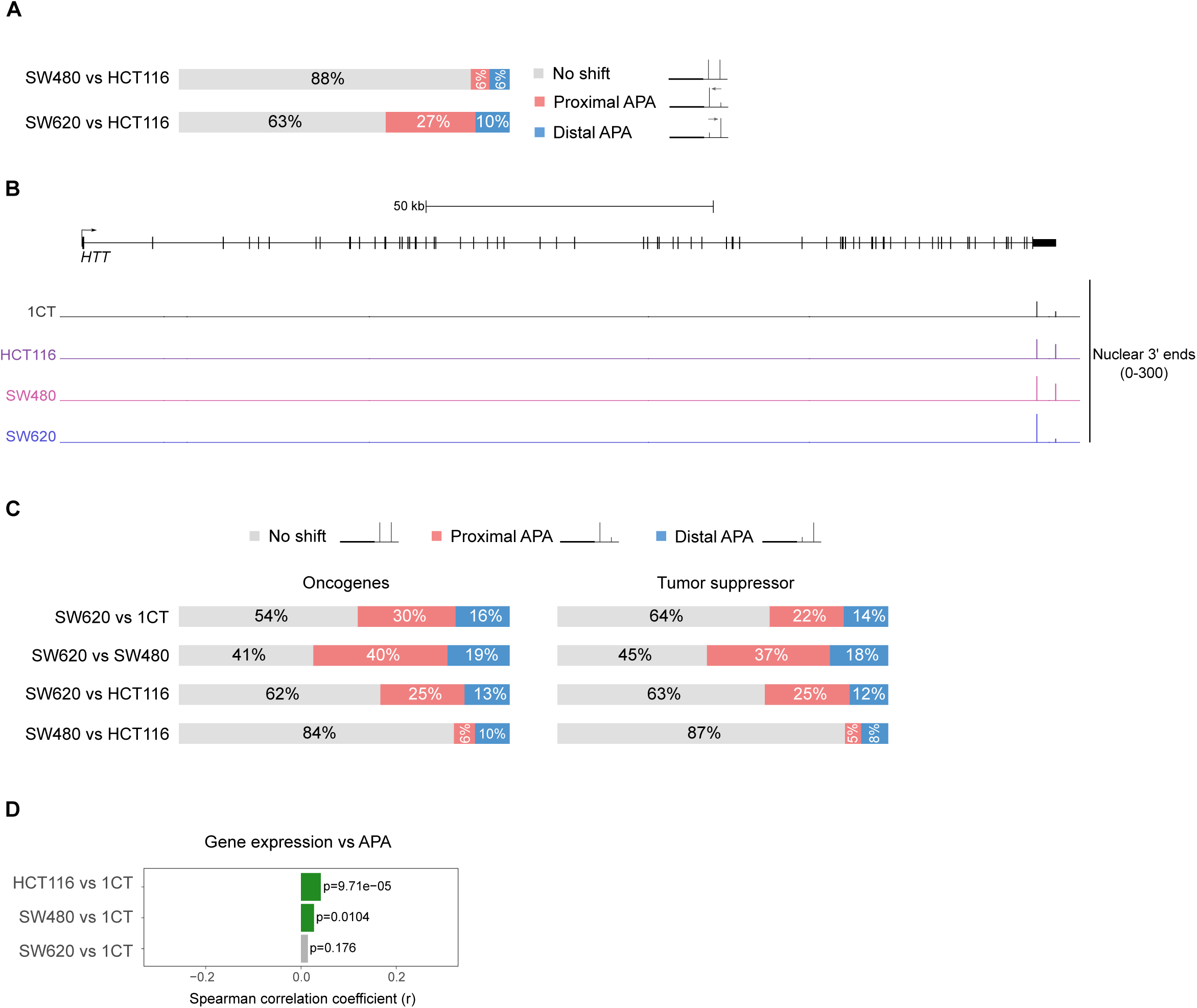
Changes in PAS selection and gene expression in the CRC progression model. Related to Figure 3. (A) Bar graph representing the alternative polyadenylation (APA) analysis in the CRC progression model. (B) Genomic profile of *HTT*, showing nuclear 3’ mRNA-seq signals. (C) Bar graph representing the APA analysis in the CRC progression model for separated oncogenes (n=533) and tumor suppressor (n=764). (D) Pairwise correlation between changes in gene expression and PAS usage (APA) in cancer cells compared to 1CT cells. Each bar represents a pairwise comparison, the bar width showing the Spearman correlation coefficient. Green bars are used for positive correlations and grey bar for non-significant correlation. Positive values indicate higher expression / distal APA, negative values lower expression / proximal APA.

**Figure S4.**
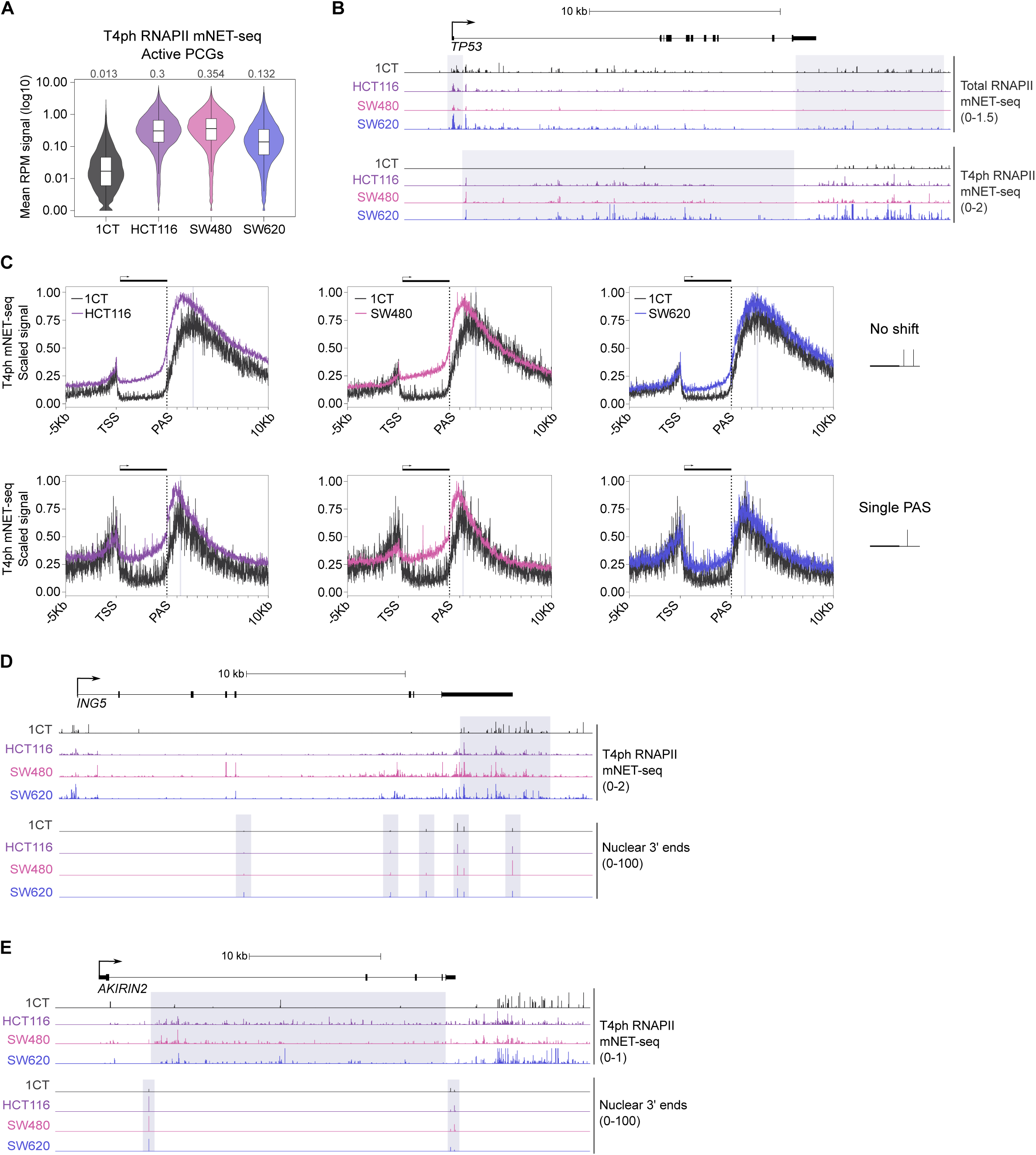
Increase in premature transcription termination in CRC; uncoupling of changes in pre-mRNA 3’ end cleavage and transcription termination during CRC progression. Related to Figure 4. (A) Violin plot representation of premature termination signal, measured by T4ph RNAPII mNET-seq across the gene body of active PCGs (n=13577). Premature termination increases strongly in cells from primary CRC, than decreases at transition to metastatic cell, yet remains upregulated compared to 1CT. (B) Genomic profile of the tumor suppressor gene *TP53*, showing increased premature transcription termination of the gene in CRC cells. Grey highlights indicate regions of TSS-associated and full-length RNAPII pausing in 1CT cells (top 4 tracks, total RNAPII mNET-seq), and premature transcription termination region in CRC cells (bottom tracks, T4ph RNAPII mNET-seq). (C) Metagene profiles of T4ph-mNET-seq signal on genes having multiple PAS but not undergoing APA (no shift, upper panel) or genes with a single PAS in the CRC progression model (lower panel). The dotted line indicates the PAS position, and the grey highlight indicates the T4ph-mNET-seq signal maximum downstream of PAS in 1CT cells. (D-E) Genomic profiles of *ING5* (D) *AKIRIN2* (E), showing T4ph-mNET-seq and nuclear 3’ mRNA-seq signals in the CRC progression model. The grey highlight indicates full-length RNAPII (D) or premature (E) termination and PAS usage.

**Figure S5.**
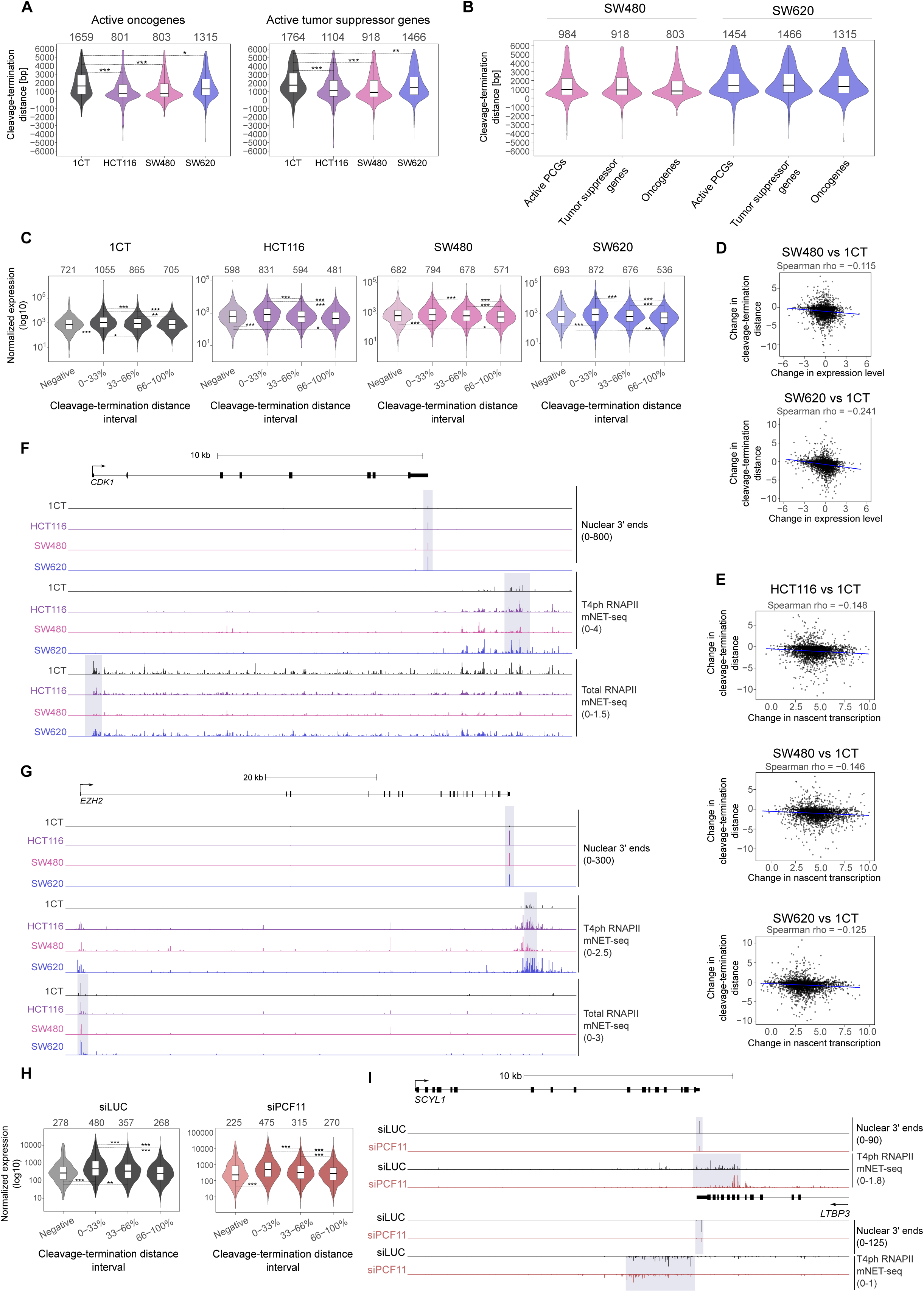
Closer proximity of PAS and the transcription termination window predicts enhanced gene expression. Related to Figure 5. (A) Violin plot representation of the cleavage-termination distance for active oncogenes (left) and tumor suppressor genes (right). (A-C, H) ***, Padj < 0.001; ** < 0.01; * <0.05. Non-indicated differences are not significant. (B) Violin plot representation of the cleavage-termination distance for active PCGs, tumor suppressor genes, and oncogenes in 1CT and SW620 cells. (C) Violin plot representation of normalized expression of genes with negative or positive cleavage-termination distance. The subset of genes with negative cleavage-termination distance was ∼ 7% of genes for 1CT and SW620; and ∼11% for HCT116 and SW480. Genes with a positive cleavage-termination distance were divided into 0-33%, 33-66%, and 66-100% distance intervals (from shortest to longest distance). (D) Scatter plots showing pairwise comparison between change in gene expression levels and change in cleavage-termination distance in SW480 or SW620 cells relative to 1CT cells. (D-E) Positive values indicate increase in expression, transcription and/or cleavage-termination distance, negative values correspond to decrease. (E) Scatter plots showing the pairwise comparison between change in nascent transcription and change in cleavage-termination distance in CRC cells relative to 1CT cells. (F-G) Genomic profile of oncogenes *CDK1* (F) and *EZH2* (G) showing nuclear 3’ mRNA-seq, T4ph-mNET-seq and Total-mNET-seq signals. Grey highlights indicate PAS usage and RNAPII pausing in 1CT cells. (H) Violin plots representing normalized expression of genes with negative (∼6% in siLUC control condition; ∼ 2.5% in siPCF11 knock-down condition) or positive cleavage-termination distance in HeLa cells. The latter group was divided into 0-33%, 33-66%, and 66-100% distance intervals. (I) Genomic profile of the convergent genes *SCYL1* and *LTBP3* locus showing nuclear 3’ mRNA-seq and T4ph-mNET-seq signals. The grey highlight indicates PAS usage and RNAPII pausing in control cells (siLUC).

